# Green Synthesized *Magnolia alba* Silver Nanoparticles Kill Pathogens, Inhibit Cancer, and Display Antioxidant and Photocatalytic Properties

**DOI:** 10.1101/2024.12.27.630567

**Authors:** Shaveen De Mel, Juliana Gruenler, Logan Khoury, Ashton Heynes, Julianne Frazekas, Kendra Damaske, Thushara Galbadage, Richard S. Gunasekera, Ross S. Anderson

## Abstract

The biosynthesis of silver nanoparticles has recently emerged as a promising approach in nanomedicine, particularly for targeted therapeutic applications. Green synthesized (plant-based) nanoparticles have been shown to offer enhanced reduction efficiency, greater bioavailability, and improved stability compared to synthetic nanoparticles. Here, we report the green synthesis of silver nanoparticles (AgNPs) using *Magnolia alba* leaf extract. The formation of these Magnolia-derived silver nanoparticles (MAgNPs) was verified through UV-Vis spectroscopy and further characterized by scanning electron microscopy (SEM) which showed that the MAgNPs have a mean diameter of 40 nm and a spherical morphology. The antibacterial efficacy of MAgNPs, evaluated by the well diffusion method, showed significant activity against *E. coli*, *Klebsiella pneumoniae*, *Pseudomonas aeruginosa*, *Enterococcus faecalis,* and strains of methicillin-resistant and -sensitive *Staphylococcus aureus.* Disc diffusion and plaque assays demonstrated notable antifungal activity against *Candida albicans* and antiviral activity against bacteriophage T7. *In vitro* studies with HCT-116 human colon cancer cells, MAgNPs exhibited significant bi-phasic inhibition of cancer cell growth. These effects were greater than that of the magnolia leaf extracts alone, confirming the green synthesized nanoparticles’ bioactive efficacy. These findings suggest that MAgNPs possess significant antimicrobial and anticancer activities, indicating their potential as therapeutic agents for certain infections and cancers. Antioxidant assays indicated that MAgNPs exhibit greater antioxidant activity than magnolia leaf extract alone. Results suggest that MAgNPs may have promise as antioxidants for treating free radical-induced disorders. Additionally, MAgNPs showed efficient photocatalytic degradation of the azo bond in methyl orange within 30 minutes, suggesting they may provide a sustainable approach to certain types of environmental pollution. To our knowledge, this is the first report of the biosynthesis of silver nanoparticles using *Magnolia alba* and examination of their antioxidant and photocatalytic properties, their killing and inhibitory effects on various bacteria, fungi, bacteriophages, and colon cancer.

## 1. Introduction

The emergence of infectious diseases and their rapid spread through pandemics have driven researchers to search for more effective medications. Traditional synthetic drugs face challenges with microbial resistance, rendering many ineffective over time. Additionally, synthetic drugs often have side effects, causing researchers to seek alternative treatments with fewer adverse effects. Attention has shifted toward natural medicines with their potentially safer profiles. Similarly, treatment of cancer is also moving from chemotherapies to targeted, green synthesized nanomedicines that have fewer side effects. As such, nanotechnology, manipulating matter at the nanoscale (1-100 nanometers), has emerged and is showing promise in the design of new, highly targeted therapies. Nanoparticles, which have unique quantum effects and high surface-to-volume ratios, have garnered interest across various scientific and industrial fields. These properties make nanoparticles useful in areas such as medicine, drug development, cancer treatment, water purification, and textile production (Wu et al., 2020).

Metallic nanoparticles exhibit distinctive surface plasmon resonance, differing from their bulk counterparts, allowing them to interact with various functional groups on polypeptides and nucleic acids. Silver nanoparticles have demonstrated broad antimicrobial properties, as well as wound healing and anti-inflammatory effects (Yaqoob et al., 2020). Synthesis of nanoparticles may be accomplished using physical, chemical, or biological methods, each having its advantages and limitations (Singh et al., 2020). Currently, the biological synthesis of AgNPs is gaining favor due to its eco-friendliness, lower toxicity, and cost-effectiveness compared to physical methods, which are energy-intensive and costly. Chemical synthesis, while efficient and relatively inexpensive, is time-consuming, and uses volatile chemicals, posing environmental risks. Researchers are therefore exploring the production of AgNPs using plants and microbes. Although microbial methods have some limitations, such as longer synthesis times and susceptibility to contamination, they offer an alternative route (Rashid et al., 2017).

Green synthesized silver nanoparticles benefit from natural stabilizers, such as phenols, flavonoids, and other phytochemicals, which stabilize the particles in suspension. In the synthesis process, phytochemicals from plant extracts reduce silver ions, forming colloidal silver particles that are stabilized by the natural compounds in the plants. This biosynthesis route is safer and avoids the need for toxic chemicals (Rajeshkumar & Bharath, 2017).

Magnolia, a plant used in traditional Asian medicine for centuries, is one such promising source for the green synthesis of AgNPs. It is valued for treating various ailments, including headaches, coughs, ulcers, and skin diseases. Magnolia’s essential oils are rich in bioactive compounds like magnolol, honokiol, linalool, α-terpineol, phenylethyl alcohol, β-pinene, and flavonoids, which contribute antioxidant, antimicrobial, anti-inflammatory, and wound-healing properties (Cheng et al., 2022). Although abundant in Asia, magnolia remains underutilized, and the plant’s full potential for medicinal and economic uses has yet to be realized.

An emerging area of interest is the role of antioxidants in combating free radicals— unstable molecules with unpaired electrons that, when present in excess, cause oxidative stress and damage cellular structures, contributing to chronic diseases like cancer, neurological disorders, and cardiovascular conditions (Phaniendra et al., 2015). Antioxidants counteract this damage by donating electrons to stabilize the free radicals (Salehi et al., 2018). The body naturally produces antioxidants, however in the event of an antioxidant deficit, synthetic antioxidants may be used but some of these have been shown to have potential carcinogenic effects later in life (Lourenço et al., 2019). Recent research suggests natural antioxidants, such as those in flavonoids and polyphenols, may be safer than synthetic alternatives. Green synthesized silver nanoparticles have demonstrated antioxidant activity comparable to natural compounds, making them a promising alternative to synthetic antioxidants (Esther Arland & Kumar, 2024).

Traditional antibiotics remain the primary treatment for infections, but their overuse has led to antibiotic resistance, posing a serious threat to global health (Chandel & Budinger, 2013). Green synthesized silver nanoparticles offer a potential solution, especially since the particles combine with phytochemicals enhancing their antimicrobial effects and making it difficult for bacteria to develop resistance (Wang et al., 2017). Green synthesized silver nanoparticles have been shown to fight drug-resistant strains effectively and prevent biofilm formation by targeting bacterial cells through both contact and ion-mediated mechanisms (Qing et al., 2018).

Green synthesized nanoparticles are being studied for their ability to degrade organic pollutants like azo dyes (Nagar & Devra, 2019). These dyes, widely used in the textile industry, are difficult to break down and can pose environmental hazards (Ganapathy Selvam & Sivakumar, 2015). Green synthesized silver nanoparticles can help by initiating photoreactions that break down pollutants thus providing a sustainable approach to pollution control by offering advantages over traditional techniques (Comparelli et al., 2005).

This study aimed to synthesize silver nanoparticles using *Magnolia alba* leaf extracts and examine their antimicrobial, anticancer, antioxidant, and photocatalytic activities. Antimicrobial activity was studied using the well diffusion method. A variety of gram-positive and gram-negative bacteria such as *E. coli*, *Klebsiella pneumoniae*, *Pseudomonas aeruginosa*, *Enterococcus faecalis*, and strains of methicillin-resistant and -sensitive *Staphylococcus aureus.* Antifungal activity was investigated against *Candida albicans* by the disc diffusion method. Antiviral activity was examined using a T7 bacteriophage plaque assay. Anticancer activity was investigated using the HCT-116 Colon Cancer cell line. Antioxidant activity was determined using TFC, TPC, TAC, FRAP, and DPPH assays, and the photocatalytic activity was analyzed by UV-Vis and by the visual loss of color in the methyl orange dye.

## 2. Material and Methodology

### 2.1 Materials

#### 2.1.1 Instruments

Incubator (COMRAD instruments), UV spectrophotometer (Bio-Rad SmartSpec Plus), autoclave (Meditry-LSB35L-1), Hot air oven (Meditary DHA-9053A), Analytical balance (OHAUS Corp, New Jersey, USA), refrigerator, Scanning Electron Microscope (VEGA, LSH Nano ZS particle size analyzer), Fume hood (Biobase-FH1000), micropipettes (Nichipet EXII, Saitama, Japan), microplate reader (BioTek Instruments, Winooski, VT, USA).

#### 2.1.2 Reagents

Folin-Ciocalteu reagent, hydrochloric acid (HCl) (CAS-7647-01-0), aluminum chloride (AlCl_3_) (CAS-7446-70-0), sodium carbonate (Na_2_CO_3_) (CAS-497-19-8), silver nitrate (AgNO_3_) (CAS-7761-88-8), sulphuric acid (H_2_SO_4_) (CAS-7664-93-9), sodium acetate buffer (C_2_H_3_NaO_2_), trisodium phosphate (Na_3_PO_4_) (CAS-7601-54-9), ammonium molybdate ([NH_4_]_6_Mo_7_O_24_·4H_2_O), ferric chloride (FeCl_3_), ammonium persulfate ([NH_4_]_2_S_2_O_8_) (CAS-7727-54-0), 1,1-Diphenyl-2-picrylhydrazyl radical (DPPH) (CAS-1898-66-4), methanol (CH_3_OH) (CAS-67-56-1), glacial acetic acid (CH_3_COOH) (CAS-64-19-7), nutrient agar, Mueller-Hinton agar, Sabouraud Dextrose agar, Luria-Bertani (LB) agar, 2,4,6-Tris(2-pyridyl)-s-triazine (TPTZ) (C_18_H_12_N_6_) (CAS-3682-35-7), sodium borohydride (NaBH_4_) (CAS-16940-66-2) Trypsin EDTA, Calf Bovine Serum, 0.1% crystal violet solution, saline solution, and distilled water.

### 2.2 Methodology

#### 2.2.1 Sample collection

*Magnolia alba* leaves were collected from the greenhouse of the Department of Biological and Physical Sciences, at The Master’s University.

#### 2.2.2 Preparation of aqueous Magnolia leaf extracts

Leaves were washed, cleaned, and dried for 2 days at 37° C. Leaves were ground to a fine powder. Five milligrams of the powdered leaves were added to 100 mL of double distilled water and incubated at 60° C for 30 minutes. Extracts were cooled and filtered using Whatman No. 1 filter paper, from each aqueous extract 1:15 dilutions were made and stored at 4° C for further analysis (Lee et al., 2013).

#### 2.2.3 Synthesis of Magnolia Silver Nanoparticles

One milliliter of leaf extracts was mixed with 9 mL of 10 mM AgNO_3_ solution. Samples were incubated at three different temperatures, 25° C (RT), 60° C, and 95° C incubated for 15 minutes, 30 minutes, 45 minutes, or 1 hour. Solutions that were at RT were maintained for 24 hours. After the incubation, 0.5 mL of each sample was diluted with 1 mL of distilled water. The absorbance was scanned between 320-520 nm by UV-visible spectrophotometry using a 10 mM AgNO_3_ solution as a reference. A 1:15 dilution of each sample was made and stored at 4° C for further analysis (Balasubramanian et al., 2015).

#### 2.2.4 Scanning Electron Microscope Statistical and Particle Size Analysis

Ten milliliters of MAgNPs were centrifuged at 4000 rpm for 10 minutes then transferred to a watch glass and dried completely at 180° C. The dried material was dissolved in 0.2 mL of distilled water and transferred into a microcentrifuge tube. (Kandiah & Chandrasekaran, 2021) To prevent degradation by light, samples were wrapped in aluminum foil and prepared for SEM analysis. The morphology and average size of the MAgNPs were determined using a VEGA LSH Nano ZS particle size analyzer at Loma Linda University, Loma Linda, CA.

#### 2.2.5 Determination of Antibacterial Activity

Antibacterial properties of MAgNPs were evaluated against several pathogenic bacteria strains: *Escherichia coli*, *Klebsiella pneumoniae*, *Pseudomonas aeruginosa, Enterococcus faecalis*, Methicillin Resistant *Staphylococcus aureus* (MRSA) and Methicillin Sensitive *Staphylococcus aureus* (MSSA) using the well diffusion method on Mueller-Hinton agar plates. Plates were inoculated with bacteria using a cotton swab under aseptic techniques. Erythromycin discs served as the positive control. The plates were incubated at 37° C for 24 hours. The zone of inhibition was then measured (Kandiah & Chandrasekaran, 2021).

#### 2.2.6 Determination of MIC and MBC_99_

To evaluate the antibacterial properties of the MAgNPs on Methicillin-resistant *Staphylococcus aureus* (MRSA) and Methicillin-sensitive *Staphylococcus aureus* (MSSA), a serial dilution minimum inhibitory concentration (MIC) assay was performed in 96-well plates. Bacterial cultures were prepared in Mueller-Hinton broth and 0.110 mL of primary culture was added to each well at an OD of 0.002. Serial dilutions were made for MAgNPs from a concentration of 4 µM and diluted to 0.008 µM. The assay was repeated with a starting concentration of 1µM and diluted to a concentration of 0.002 µM. An equivalent concentration of MLE was also used as a control. In assays, a bacteria-only positive control and a media-only negative control were used. After incubation, samples from the dilution assay were spot-plated to determine the MIC and the minimum bactericidal concentration (MBC_99_) of the MAgNPs. The MIC of the MAgNPs is the lowest concentration at which no bacterial growth was observed. The MBC_99_ is the lowest concentration of MAgNP at which 99% of the bacteria were killed (Andrews, 2001).

#### 2.2.7 Determination of Antifungal activity

The antifungal properties of MAgNPs were evaluated against the pathogenic fungus *Candida albicans* using the disc diffusion method on Sabouraud Dextrose Agar plates. The fungus was evenly inoculated on the plates using a cotton swab under aseptic conditions. Sterile water served as the negative control, and Fluconazole (a gift of Dr. John Roueche) was used as the positive control. The test samples, leaf extract, and MAgNPs were added to discs. The plates were incubated at 37° C for 48 hours. The resulting zone of inhibition was then measured (Taufikurohmah & Tantyani, 2020).

#### 2.2.8 Determination of Antiviral activity

To test the antiviral properties of MAgNP’s against bacteriophages, T7 coliphage (bacteriophage) which targets *E. coli* BL21 stain was used. The antiviral activity of MAgNP was quantified through a plaque count assay. The bacteriophages with a stock concentration of ∼1×10^10^ PFU/mL were first serially diluted to 1×10^-3^, exposed to MLE (50 mg/l), and MAgNP (0.6 mM), and incubated for 20 minutes at room temperature. After incubation, the treated bacteriophages were further diluted 1×10^-5^ to ensure the plaques were reliably countable. The diluted bacteriophages were mixed well with a fresh culture of *E. coli* BL21 (OD 0.02) by vortexing, and mixed with warmed LB top agar, mixed well again by vortexing, and poured onto LB agar plates. The plates were then incubated overnight at 37° C. The plaques were counted, and antiviral activity was calculated.

#### 2.2.9 Determination of Anticancer Properties

##### 2.2.9.1 Cell Culture

Colorectal cancer cell line, (HCT-116 was cultured in Dulbecco’s Modified Eagle Medium supplemented with 10% Calf Bovine Serum at 37° C in a 5% CO_2_ environment. At 85-90% confluence the cells were treated with 0.25% Trypsin EDTA and brought into suspension. Cells were seeded in 96-well plates with a density of 0.01 x 10^6^ cells/well and allowed to adhere overnight before treatment.

##### 2.2.9.2 Crystal Violet Cell Proliferation Assay

HCT-116 cells initially plated in 96-well plates, were treated with various concentrations of MAgNPs for 48 hours. Similarly, various concentrations of magnolia leaf extract treatment alone and silver nitrate treatments alone were also tested. Following treatments, the cells were exposed to a 0.1% crystal violet solution for 30 minutes at room temperature and rinsed with deionized water to remove excess dye. Absorbance readings at 570 nm were taken using a microplate reader.

#### 2.2.10 Determination of Antioxidant Properties

##### 2.2.10.1 Total Flavonoid Content

Total flavonoid content (TFC) was estimated as per the aluminum chloride colorimetric method (Kumaresan et al., 2019). A 1 mL sample of extract was added to 3 mL of 5% NaNO_3_, mixed, and kept for 1 minute. A 0.1 mL aliquot of 10% AlCl_3_ was added and incubated for 5 minutes at RT after which 0.5 mL of 1M NaOH was added. The absorbance was measured at 415 nm with distilled water as a blank. The results are expressed in μg of Quercetin equivalents per 100 g (Kumaresan et al., 2019).

##### 2.2.10.2 Total Phenolic Content

Total phenolic content was estimated using the Folin-Ciocalteu colorimetric method. A 1.5 mL sample of leaf extract was mixed with 0.1 mL of FC reagent and mixed for 5 minutes after which 0.2 mL of 20% Na_2_CO_3_ was added, to both the leaf extract and the MAgNP. Samples were prepared in triplicate and incubated for 1 hour in the dark at RT. The absorbance was measured at 765 nm using distilled water as a blank. The results are expressed in grams of Gallic acid equivalents per 100 g (Kumaresan et al., 2019).

##### 2.2.10.3 Total Antioxidant Capacity

Total antioxidant capacity was estimated using the phosphomolybdenum assay (Perera & Kandiah, 2018). A sample extract of 1.5 mL was mixed with 0.5 mL phosphomolybdenum reagent (0.6 M sulphuric acid, 4 mM ammonium molybdate, 28 mM sodium sulfate in 1:1:1 ratio), afterwards, the samples were incubated at 90° C for 90 minutes and cooled. The absorbance was read at 695 nm using distilled water as a blank. The results are expressed in grams of ascorbic acid equivalents per 100 g.

##### 2.2.10.4 Determination of Ferric Reducing Antioxidant Power (FRAP)

The colorimetric method was used to determine the ferric-reducing ability of the leaf extracts (Balasubramanian et al., 2015). The FRAP reagent was created by adding 50 mL of a 0.3 M acetate buffer, pH 3.6, containing 5 mL of 10 mM TPTZ to a solution of 40 mM HCl containing 5 mL of 20 mM FeCl_3_·6H_2_O. A 0.1 mL aliquot of leaf extract or MAgNP was added to 2.9 mL of the FRAP reagent and the absorbance was recorded at 1-minute intervals at a wavelength of 593 nm, using distilled water as a blank.

##### 2.2.10.5 Determination of 2,2-diphenyl-1-picrylhydrazyl Activity

One milliliter of either leaf extract or MAgNP was mixed with 2 mL of a 0.004% DPPH solution incubated at RT for 30 minutes. The absorbance of the mixture was measured at 517 nm, with methanol used as a reference. The following formula was used to determine the sample’s % DPPH scavenging capacity to neutralize DPPH free radicals:

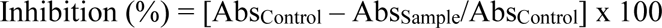

Where Abs_Control_ is the absorbance of the DPPH reagent only, and Abs_Sample_ is the absorbance of either leaf extract or MAgNP after incubation of DPPH (Perera & Kandiah, 2018).

#### 2.2.11 Phytochemical Analysis

A qualitative phytochemical analysis as described by Roghini & Vijayalakshmi was conducted on diluted MLE to detect the presence of various secondary metabolites, including carbohydrates, proteins, saponins, steroids, tannins, and terpenoids (Roghini & Vijayalakshmi, 2018).

#### 2.2.12 Determination of Photocatalytic Activity

One milliliter of 9 mM MAgNP and 1 mL of 0.2 M NaBH_4_ were added to a 100 mL solution of 2 mM Methyl Orange. The absorbance of the mixture was then measured at 5-minute intervals for 30 minutes while scanning from 300 nm to 560 nm. Distilled water was used as a blank (Kandiah & Chandrasekaran, 2021).

## 3. Results & Discussion

Silver nanoparticles (AgNPs) have distinct physiochemical properties and are employed in a wide range of applications. Green synthesis of AgNPs, in comparison to other methods of synthesis, is both a simple and rapid method. It has shown promise as it is environmentally friendly, cost-effective, sustainable, and reproducible. At the time of this writing, there has been no documented research using *Magnolia alba* leaves for the green synthesis of AgNPs.

Green synthesized silver nanoparticles have been shown to have high antioxidant and free radical scavenging properties as compared to vitamin C (Keshari et al., 2020). Green synthesized particles permit molecules such as phenols and flavonoids to bind the nanoparticle surfaces forming a stabilizing layer that confers long-term stability (Kuppusamy et al., 2016). The MAgNPs we synthesized remained stable and effective for approximately six months when stored at 4° C. Water was chosen as the extraction medium due to its non-toxicity and suitability for herbal products in comparison to other commonly used solvents such as methanol. During AgNPs synthesis, Ag^+^ is reduced to Ag^0^ which is stabilized in the aqueous extracts by biomolecules and secondary metabolites (Zhao et al., 2014).

### 3.1 MAgNP Synthesis and Characterization

#### 3.1.1 MAgNP Synthesis

Synthesis of MAgNPs was primarily detected by the reddish-brown color change in the solutions (Figure 1 A). This is due to the formation of AgNP clusters or aggregates, which absorb and scatter light in a way that gives rise to the reddish-brown color, which implies the formation of MAgNPs (Ahmed et al., 2016). As a result of the segregation of MAgNP, samples turned black over time (Cao et al., 2011). Optimizing the synthesis of AgNPs we found that prolonged incubation time and high temperature facilitated the reduction of silver ions leading to higher AgNP concentrations. This process, aided by Brownian Motion (continuous bombardment between silver ions and phytochemicals), resulted in the aggregation of silver and phytochemicals through the LaMer mechanism (Piñero et al., 2017). Synthesis of MAgNPs was further confirmed by UV-Vis spectroscopy scanning between 320 and 520 nm. Green synthesized silver nanoparticles are expected to exhibit a peak absorbance between 420 and 450 nm, we observed a peak at 440 nm (Figure 1B).

**Figure 1.**
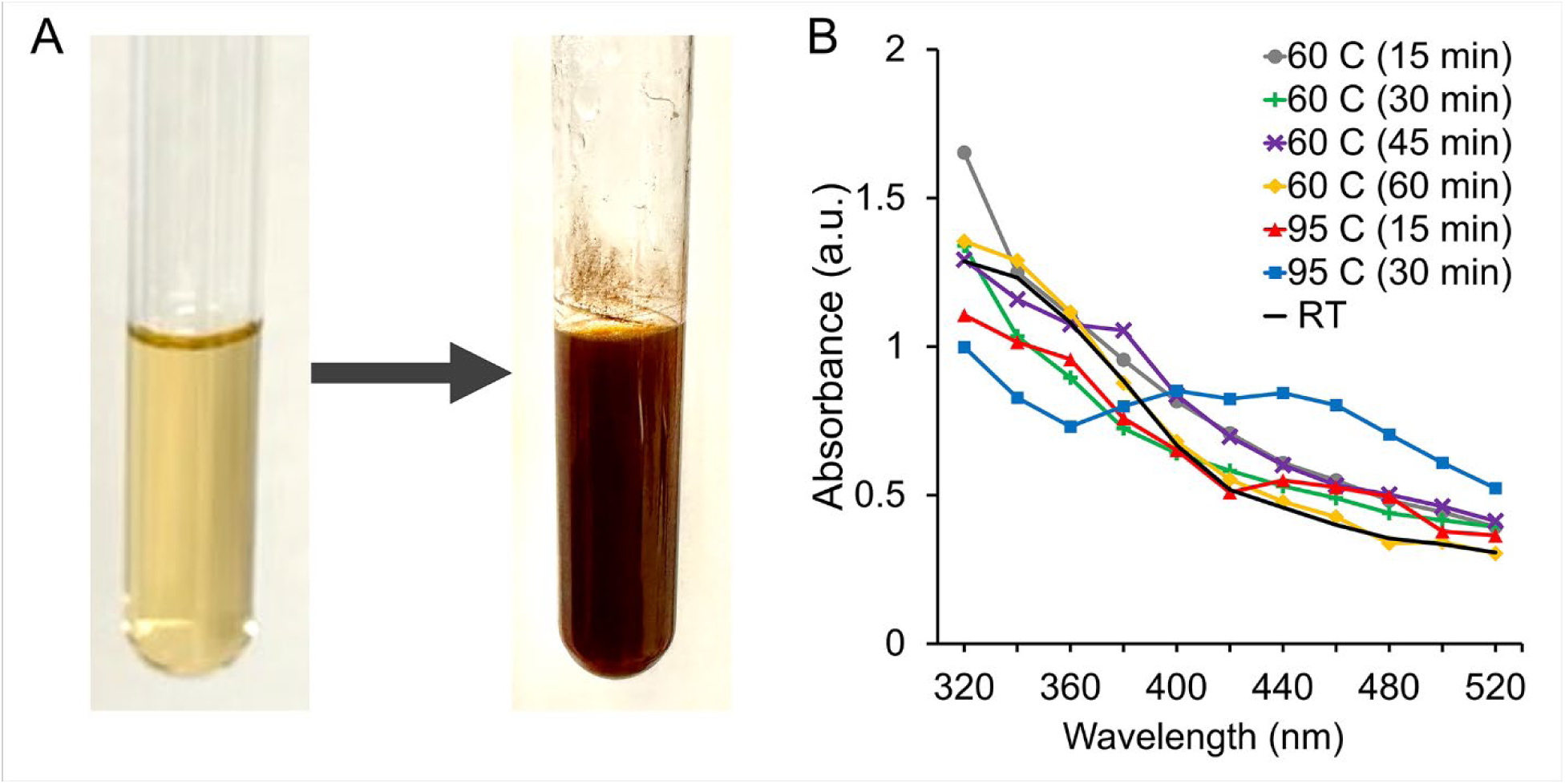
Green synthesis and Uv-Vis characterization of MAgNPs. (A) Color change observed before and after MAgNPs synthesis. (B) The spectroscopic analysis of the MAgNPs synthesized for different periods and at different temperatures. Note the peak absorbance at 440 nm at 95° C at 15 and 30 minutes which is indicative of the presence of silver nanoparticles. There was no further change in peak height at 45 or 60 minutes.

Temperature is a significant factor in the green synthesis of AgNPs. Synthesis of MAgNPs was optimal when the temperature was maintained at 95° C for 30 minutes. Experiments carried out at 25° C and 60° C failed to create MAgNPs. This is consistent with other findings (Lee et al., 2013). The higher temperature could reduce the time required for bio-reduction thus accelerating the formation and growth of AgNPs resulting in greater concentrations. This is consistent with the idea that with Brownian Motion and the LaMer mechanism such conditions enhance aggregation of Ag^+^ with phytochemicals (Darroudi et al., 2011; Piñero et al., 2017).

#### 3.1.2 Scanning Electron Microscopic (SEM) Images of MAgNPs

The morphology of the AgNPs was examined using Scanning Electron Microscopy (SEM) (Figure 2). SEM images revealed that the particles were spherical and had a mean diameter of 40 nm. Similar results were observed by Jain, A.S. et al. (Jain et al., 2021).

**Figure 2.**
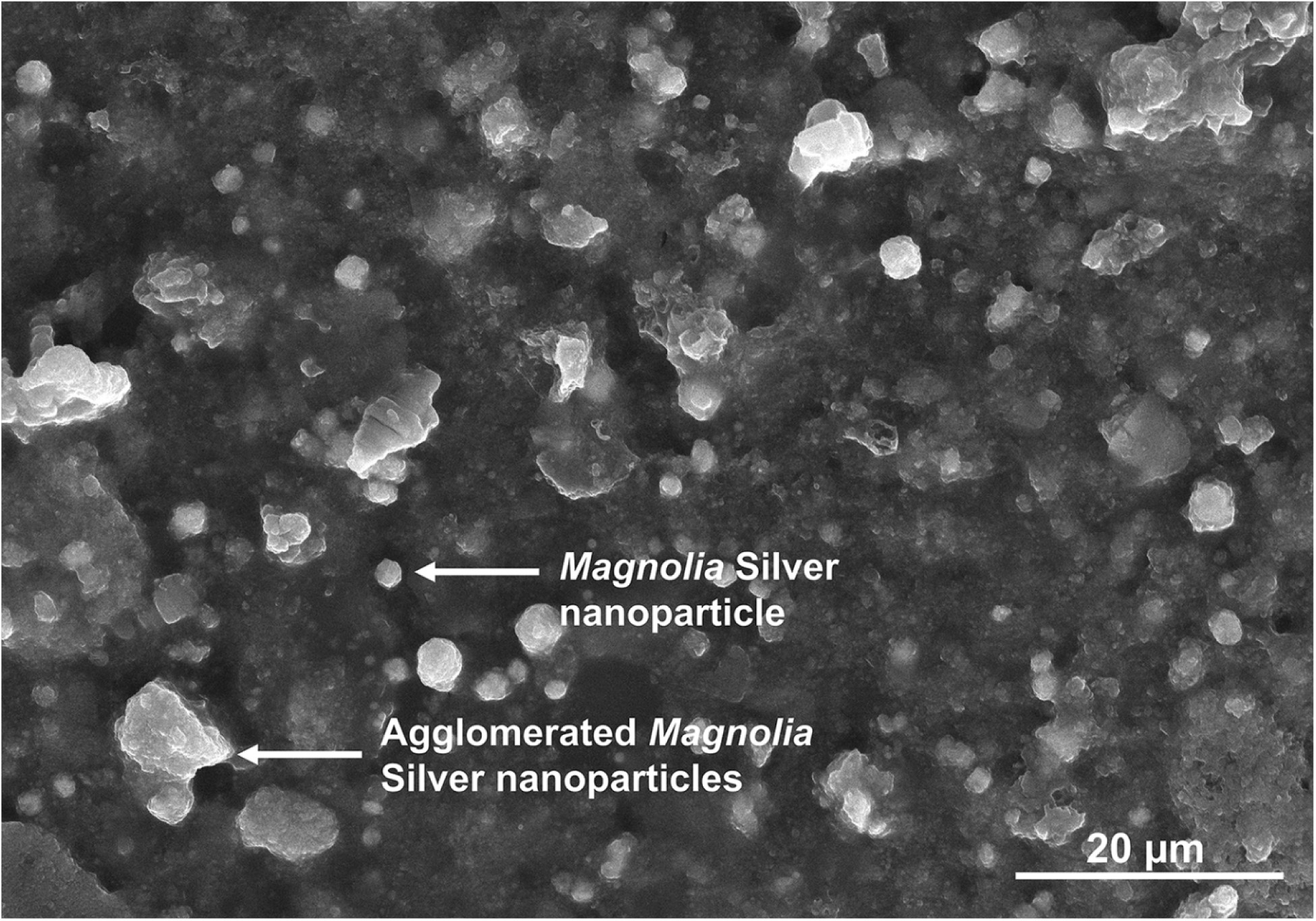
Characterization of MAgNPs using scanning electron microscopy (SEM) at 30.0 Kv. SEM revealed that *Magnolia* silver nanoparticles (MAgNPs) had a mean diameter of 40 nm and were spherical. Agglomerated *Magnolia* silver nanoparticles were also observed as irregularly shaped clumps.

### 3.3 Determination of Antibacterial Activity

Silver nanoparticles have a built-in ability to fight microbes, which is boosted by their high surface area allowing them to exhibit a broad-spectrum antimicrobial effect. Y. Qing et al. have proposed a mechanism for how AgNPs work (Qing et al., 2018). The AgNPs initially attach to the bacterial cell wall, disrupting it, and causing the cell’s contents to leak out, ultimately leading to death. Additionally, silver ions can bind to proteins essential for energy production and prevent the generation of ATP. Inside the bacterial cell, silver ions react with biomolecules, generating ROS that induce apoptosis, or cell death, through both contact killing and ion-mediated killing (Qing et al., 2018).

This study investigated the antibacterial properties of MAgNPs against various bacterial strains. Well diffusion assays revealed significant antibacterial activity against *E. coli*, *K. pneumoniae*, *P. aeruginosa*, *E. faecalis*, MSSA, and MRSA (Figure 3 A to F). The presence of a clear zone of inhibition (ZOI) around wells containing 0.6 mM MAgNPs indicates antibacterial activity (Table 1). According to Muharni’s criteria, ZOI diameters were categorized as follows: weak (<10 mm), moderate (10-15 mm), and strong (>15 mm) (Taufikurohmah & Tantyani, 2020). Based on this categorization, *E. coli*, *P. aeruginosa*, *E. faecalis*, MSSA & MRSA exhibited strong antibacterial activity, with ZOIs greater than 15 mm. The ZOI for *K. pneumoniae* was found to be in the moderate category, which may be attributed to the thickness of the polysaccharide layer (∼160 nm) and unique fiber arrangement, which makes it a virulent pathogenic strain. The leaf extracts exhibited a slight, insignificant ZOI, suggesting that their phytochemicals may have some antibacterial properties. Similar results were obtained by Anbumani, D. et al., which confirms the antibacterial efficacy of the green synthesized nanoparticles (Anbumani et al., 2022).

**Figure 3.**
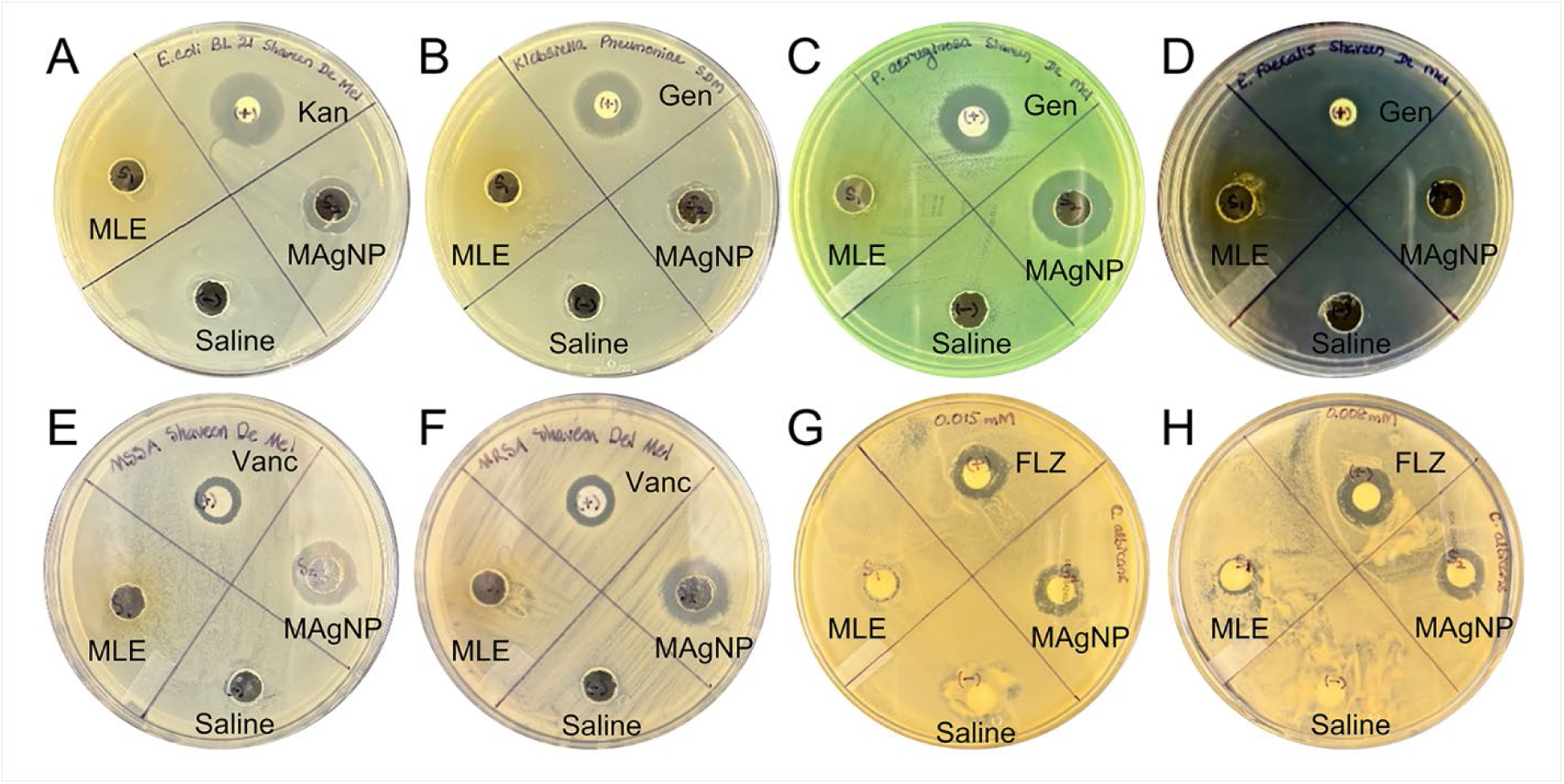
Well and disc diffusion assays for antibacterial and antifungal properties of MAgNPs against medically relevant pathogens. Kanamycin (Kan, 100 μg/mL), Gentamycin (Gen, 50 μg/mL), Vancomycin (Vanc, 50 μg/mL), Fluconazole (FLZ 50 mM) Magnolia Silver Nanoparticles (MAgNP, 0.6 mM), Magnolia leaf extract (MLE), and Saline were exposed to a lawn of bacteria and grown at 37° C for 24 hours. (A) *E. coli* BL21, (B) *Klebsiella pneumoniae*, (C) *Pseudomonas aeruginosa*, (D) *Enterococcus faecalis,* (E) MSSA, (F) MRSA and (G and H) *C. albicans.* (G) 0.015 mM of MAgNPs and (H) 8 μM of MAgNPs.

**Table 1.** Zone of inhibition for each of the treatments in the well-diffusion assay.

| Bacterial species<sup>1</sup> | Positive Control (mm)<sup>2</sup> | MAGNP<sup>3</sup> (0.6 mM) (mm) | Magnolia Leaf Extract (mm) | Negative Control, Saline (mm) |
| --- | --- | --- | --- | --- |
| *E. coli* | 18 | 16 | 0 | 0 |
| *K. pneumoniae* | 14 | 14 | 0 | 0 |
| *P. aeruginosa* | 18 | 19 | 0 | 0 |
| *E. faecalis* | 12 | 16 | 0 | 0 |
| MSSA | 11 | 16 | 0 | 0 |
| MRSA | 12 | 16 | 0 | 0 |
| *C. albicans* | 15 | 14* | 0 | 0 |
|  | 15 | 14** | 0 | 0 |

<sup>1</sup> Bacterial strain information: *Escherichia coli* BL 21; *Klebsiella pneumoniae* ATCC# BAA-1144; *Enterococcus faecalis* ATCC# 51299; *Pseudomonas aeruginosa* ATCC# 27853; Methicillin Sensitive *Staphylococcus aureus* (MSSA) ATCC# 25923; Methicillin Resistant *Staphylococcus aureus* (MRSA) ATCC# 33591, *Candida albicans* ATCC# 10231
<sup>2</sup> Zone of inhibition measured in mm. Positive controls: *E. coli* – kanamycin; *K. pneumoniae*, *P. aeruginosa*, and *E. faecalis* – gentamicin; MSSA and MRSA - vancomycin; *C. albicans* - fluconazole
<sup>3</sup> Magnolia silver nanoparticles (MAGNP)
\* MAGNP concentration 0.015 mM
\*\* MAGNP concentration 8 µM

### 3.4 Determination of MIC & MLC

MAgNPs exhibited potent antibacterial activity against both MRSA and MSSA in 96-well plate serial dilution assay, with MICs of 0.015 mM and MBC_99_ of 0.015 and 0.06 mM (Figure 4 and Table 2). Magnolia leaf extract alone demonstrated no significant antibacterial activity, confirming the enhancement of bioactivity through the synthesis of MAgNPs. These findings suggest that MAgNPs are effective in combating antibiotic-resistant pathogens such as MRSA.

**Figure 4.**
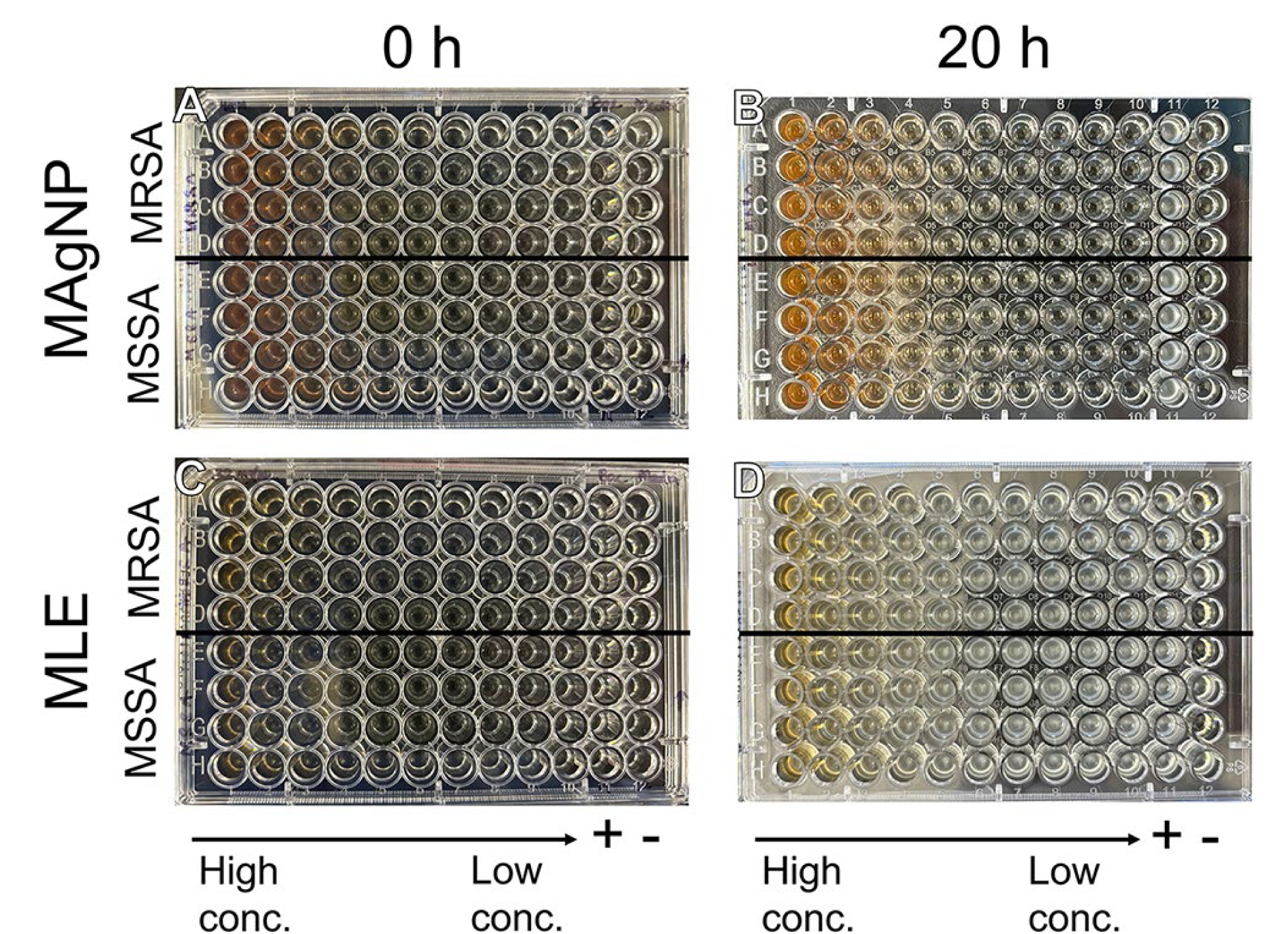
Micro dilution assay for minimal inhibitory concentration (MIC) determination. Methicillin Resistant *Staphylococcus aureus* (MRSA), Methicillin Sensitive *Staphylococcus aureus* (MSSA) was with magnolia silver nanoparticles (MAgNP) (A and B), and Magnolia leaf extract (MLE) (C and D). (A and C) before incubation, (B and D) after 20 hours of incubation. In each 96-well plate, column 11 contained bacteria only (positive control) and column 12 contained media only (negative control). (A) MRSA and MSSA treated with MAgNP using serial dilution at concentrations of 4 mM to 0.008 mM at the start of the experiment (0 h). (B) MRSA and MSSA treated with MAgNP after 20 hours of incubation at 37° C. (C) MRSA and MSSA treated with MLE using serial dilution at concentrations of 32 mg/mL to 0.06 mg/mL at the start of the experiment (0 h). (D) MRSA and MSSA treated with MLE after 20 hours of incubation at 37° C.

**Table 2.**
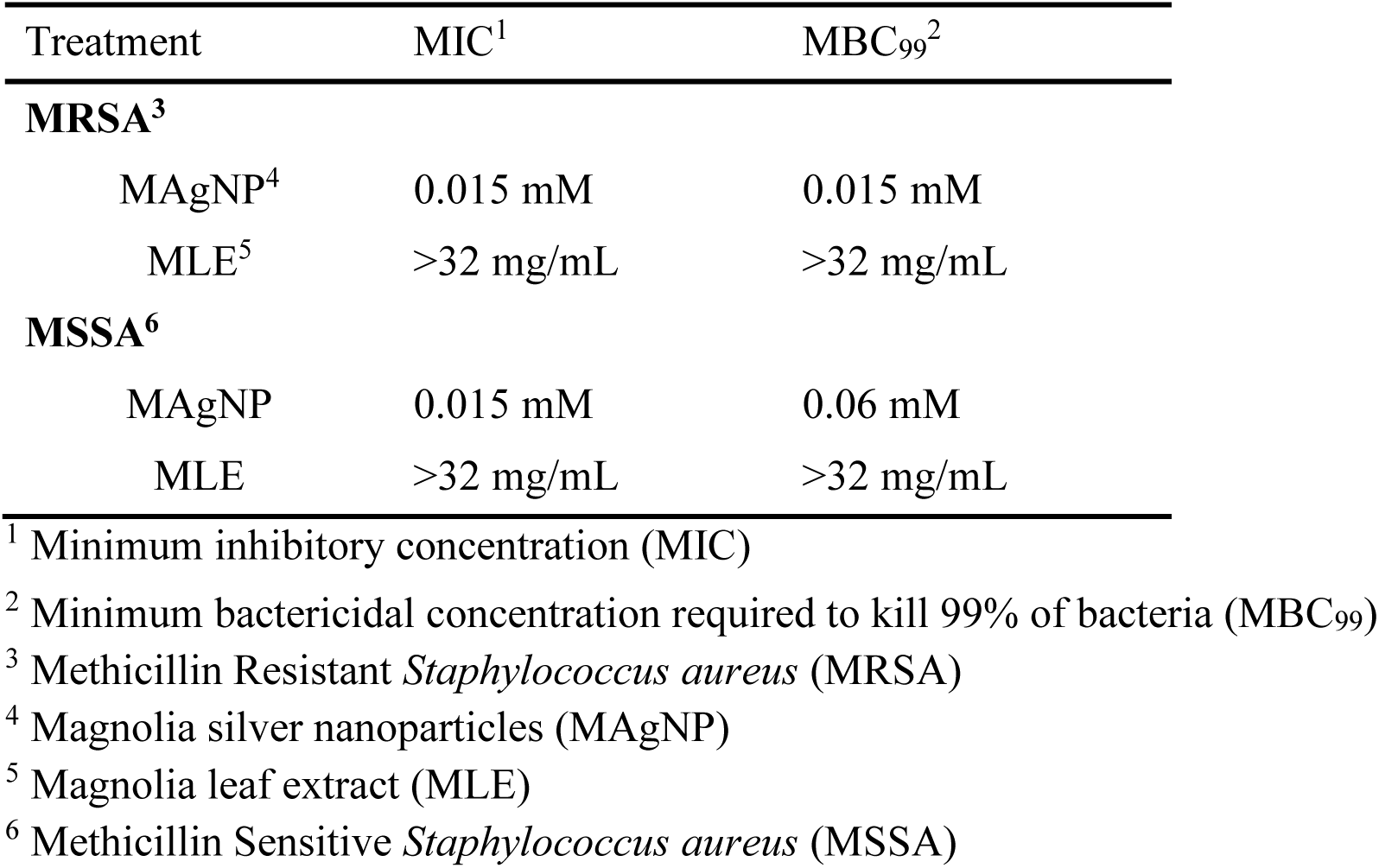
Minimum inhibitory concentration (MIC) and minimum bactericidal concentration of 99% (MBC_99_) for magnolia leaf extract (MLE) and magnolia silver nanoparticles (MAgNP)

The antibacterial activity of MAgNPs observed in this study highlights their potential as a novel therapeutic agent against resistant bacterial strains, including MRSA. The enhanced efficacy of MAgNPs compared to MLE alone underscores the synergistic effects of green synthesis, where bioactive phytochemicals stabilize and enhance nanoparticle properties. This approach not only reduces reliance on conventional chemical antibiotics but also aligns with sustainable and eco-friendly practices in nanomedicine. The consistency of MIC and MBC_99_ results reinforces the reproducibility and reliability of MAgNPs as an antibacterial agent. The well-diffusion assay results further support this finding, showing clear zones of inhibition that exceed those of MLE alone.

However, this study has certain limitations, including the need for further exploration of cytotoxicity and the mechanisms underlying bacterial inhibition. Future research should investigate the effects of MAgNPs on biofilms and assess their efficacy in complex biological matrices to simulate clinical conditions better. The antimicrobial results against both MRSA and MSSA, with MIC values of 0.015 mM, demonstrate that MAgNPs maintain potent antibacterial activity against both antibiotic-sensitive and resistant strains. This suggests that the mechanism of action differs from conventional antibiotics, potentially offering a new strategy to combat antimicrobial resistance (Durán et al., 2016). MAgNPs likely exert their antimicrobial activity through generating ROS that induce oxidative stress, disrupting bacterial cell membrane integrity, and interfering with essential intracellular components such as proteins, enzymes, and DNA.

### 3.5 Determination of Antifungal Activity

Antifungal properties of MAgNPs were also investigated using *Candida albicans*. Triplicate experiments revealed that MAgNPs exhibit inhibitory properties, evident from the ZOI around the disc (Table 1), at concentrations of 0.015 and 0.008 mM of MAgNPs (Figure 3 G & H). According to Johannes, E. et al., antifungal activity can be categorized based on inhibition zone diameter: weak (<5 mm), moderate (5-10 mm), strong (11-20 mm), and very strong (>20 mm) (Johannes et al., 2019). Notably, MAgNPs demonstrated potent antifungal activity, as evidenced by their significant ZOI across the two concentrations used, thus categorizing them as a strong antifungal against the nosocomial (Hospital Acquired Infection) pathogenic *Candida albicans*. Kim, K. J. et al., hypothesize that MAgNPs disrupt membrane permeability by perturbing lipid bilayers, leading to ion leakage, pore formation, and dissipation of electrical potential (Kim et al., 2009). Thus, MAgNPs may inhibit *Candida albicans* growth by compromising membrane integrity. Green synthesized AgNPs exhibit strong antifungal activity against this pathogen. In this assay, the leaf extracts alone exhibited an immeasurable and insignificant ZOI, implying that their phytochemicals may have some antifungal properties. Similar results have been reported by Basem and Enas, confirming the antifungal activity of the green synthesized silver nanoparticles (Abdallah & Ali, 2022).

### 3.6 Determination of Antiviral Activity

MAgNPs significantly reduced the plaque-forming ability of T7 bacteriophages, indicating strong antiviral activity compared to a no-exposure group (p < 0.0001). MAgNPs also showed significantly better antiviral activity compared to that of MLE (p =0.0004) (Figure 5). This suggests that nanoparticles synthesized using Magnolia extracts can effectively reduce bacteriophage viability. The antiviral properties demonstrated by MAgNPs represent a significant advancement in the application of green synthesized nanoparticles for antiviral therapy. The ability to inhibit bacteriophage activity serves as a model for potential applications against human viruses, positioning MAgNPs as promising candidates for antiviral drug development.

**Figure 5.**
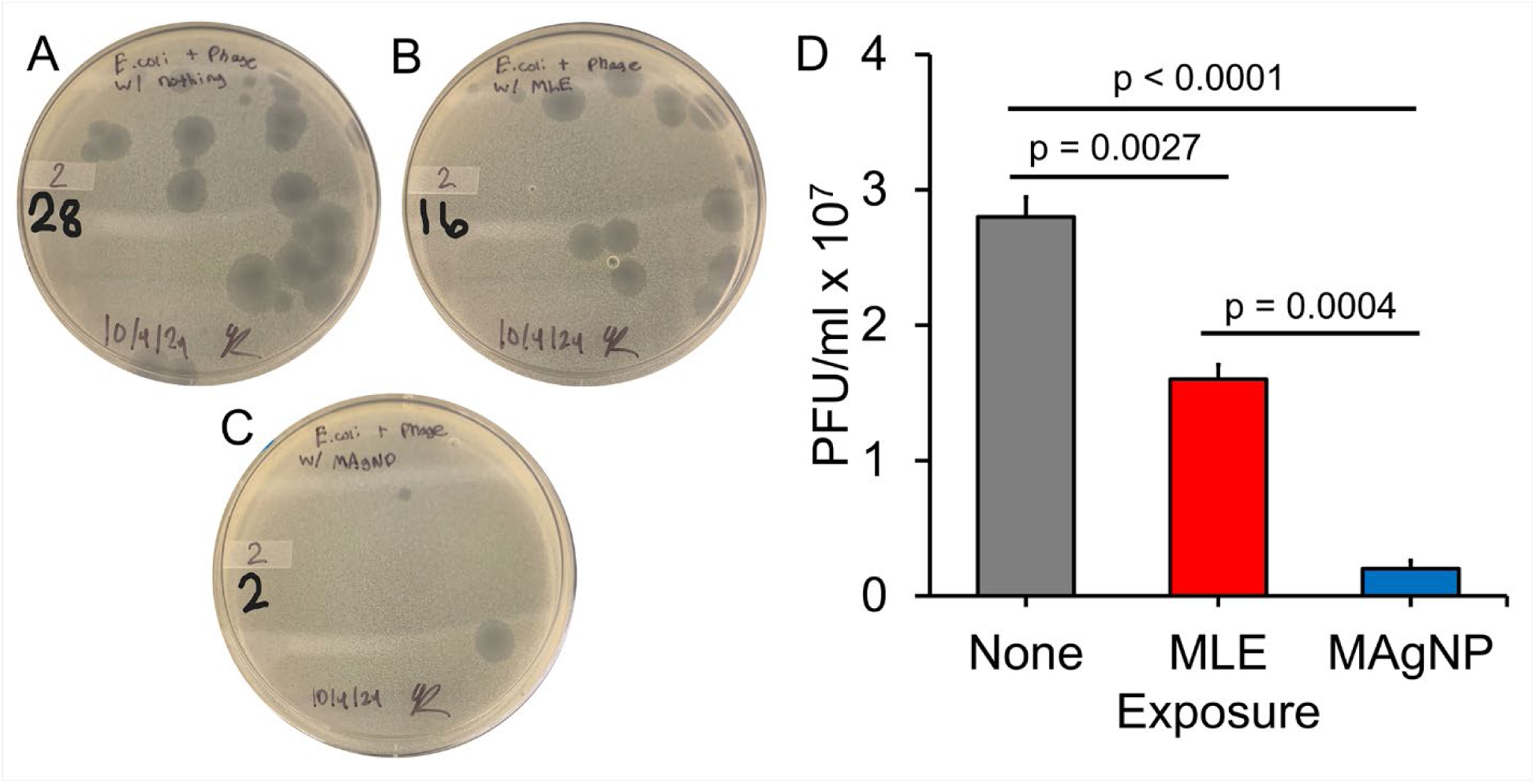
Bacteriophage plaque assay to examine the antiviral activity of Magnolia silver nanoparticles (MAgNP) T7 coliphage (bacteriophage) exposed to different agents for 20 minutes and then incubated with *E. coli* BL21 strain for plaque formation. (A) no exposure, (B) Magnolia leaf extract (MLE), and (C) Magnolia silver nanoparticles (MAgNP). (D) PFU/mL for each of the exposure groups after incubating at 37° C overnight. This experiment was repeated at least 3 times, and a representative data set is shown. The error bars are the standard error of the mean (SE). P-values were obtained using a t-test to compare the means of the groups.

The MAgNPs may bind to the viral capsid, tail, or tail fiber, causing structural damage and making the phages non-viable. These actions highlight the potential of these nanoparticles as broad-spectrum antimicrobial agents (Gilcrease et al., 2020). The findings contribute to the growing body of evidence supporting the role of silver nanoparticles as versatile antimicrobial agents and highlight the advantages of green synthesis for enhancing bioactivity while minimizing environmental impact. However, the reliance on a bacteriophage model may not fully capture the complexity of interactions in human or mammalian viral infections. While the results are promising, further studies are necessary to evaluate MAgNPs’ antiviral effects against clinically relevant viruses, elucidate their mechanisms of action, and determine their safety in host systems.

### 3.7 Anticancer Property

This biphasic trend is different from the monotonic dose-dependent response typically seen with conventional chemotherapeutic agents which is seen usually in toxicology studies (Calabrese & Baldwin, 2003). Such a dose-response trend shows a proportional increase in response to increasing concentrations of the toxic substance until a maximum cytotoxicity effect is reached. The enhanced efficacy of MAgNPs compared to MLE) alone indicates a synergistic -not additive-effect between the silver nanoparticles and the plant bioactive compounds present in *Magnolia alba* (Figure 6). This synergy likely stems from the combinatorial inherent cytotoxic properties of silver nanoparticles and bioactive molecules like magnolol and honokiol which are known anticancer compounds found in Magnolia species. These have been previously documented to possess pro-apoptotic and antiproliferative activity against various cancer cell lines (Lee et al., 2011; Park et al., 2012).

**Figure 6.**
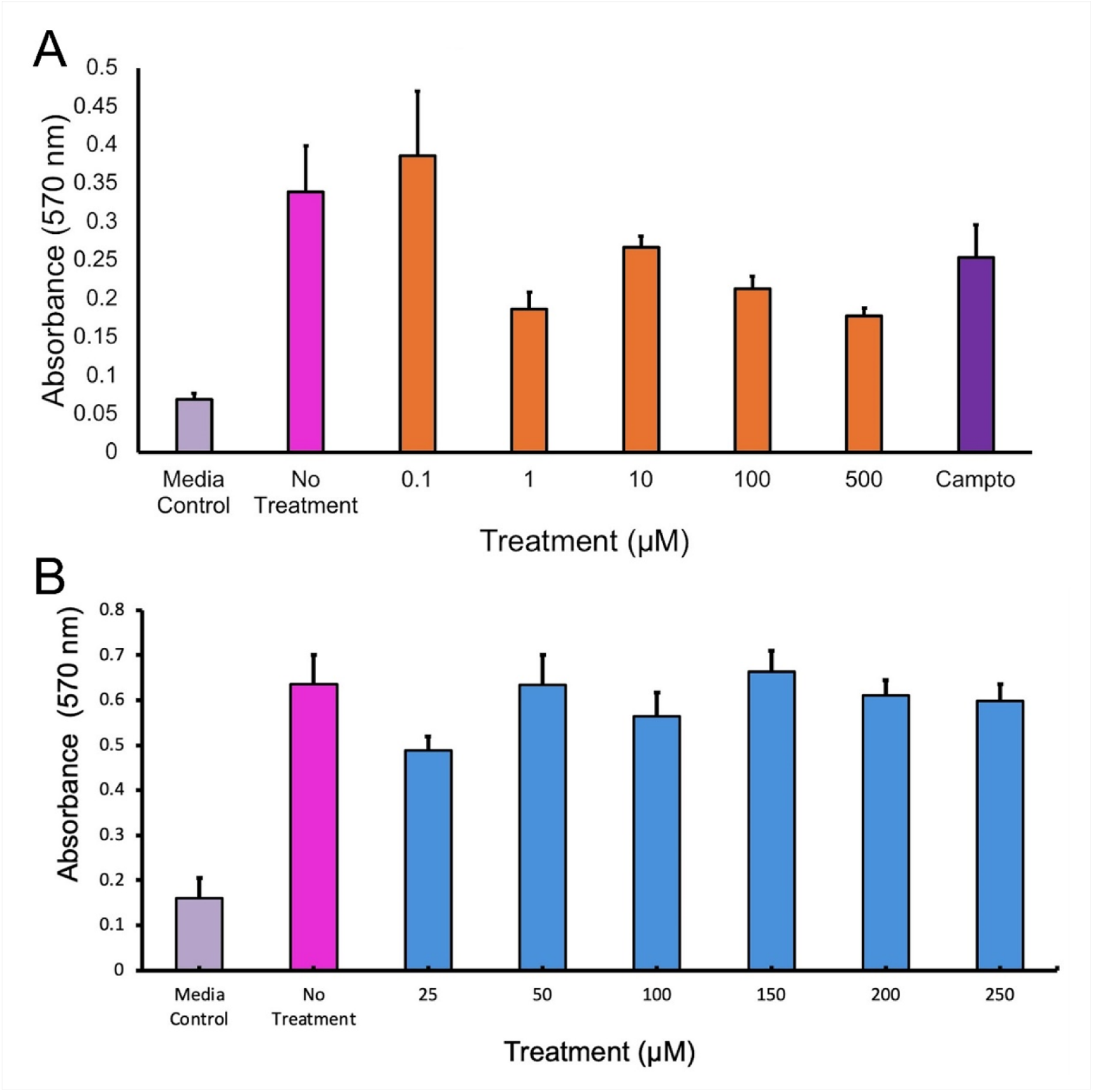
Crystal violet assay for the effects of MAgNP and magnolia leaf extract (MLE) on HCT-116 cells. (A) MAgNP concentration range, from 0.1 to 500 µM in 10-fold serial dilutions. (B) Selected MAgNP concentration range, from 25 to 250 µM. The MAgNPs depict an inhibitory bi-phasic effect on HCT-116 colorectal cancer cells. The biphasic trend showed enhanced inhibition of cells at lower concentrations and then again at higher concentrations. This suggests more than one mechanism of action which could indicate distinct cellular pathways being activated at different concentrations usually seen with antiproliferative and pro-apoptotic mechanisms of natural plant-derived bioactive molecules (PBAMS) also known as phytochemicals. The error bars are the standard error of the mean (SE).

The size of the synthesized MAgNPs (approximately 40 nm) falls within the optimal range for cellular uptake and interaction with biological systems. Previous studies have shown that the spherical morphology observed through SEM analysis contributes to optimal cellular interaction and uptake efficiency (Albanese et al., 2012). Furthermore, nanoparticles in this size range are known to effectively penetrate cancer cells while maintaining stability in biological media (Jiang et al., 2008).

### 3.8 Determination of Antioxidant Activity

#### 3.8.1 Total Flavonoid Content (TFC)

Total Flavonoid Content was estimated as per the aluminum chloride colorimetric method described by Kumaresan *et al*., (Kumaresan et al., 2019). Principle implicates aluminum chloride ions to form acid stable complexes with the C-4 keto group and either the C-3 or C-5 hydroxyl group of flavones and flavonols, hence the technique is used to evaluate flavonoids with extreme absorption at 415 nm. It also forms acid-labile complexes with the orthodihydroxyl groups in flavonoids’ A- or B-rings (Sabli et al., 2012). MAgNPs showed higher TFC values compared to that of leaf aqueous extracts (Figure 7 A), similar findings were reported previously (Yarrappagaari et al., 2020).

**Figure 7.**
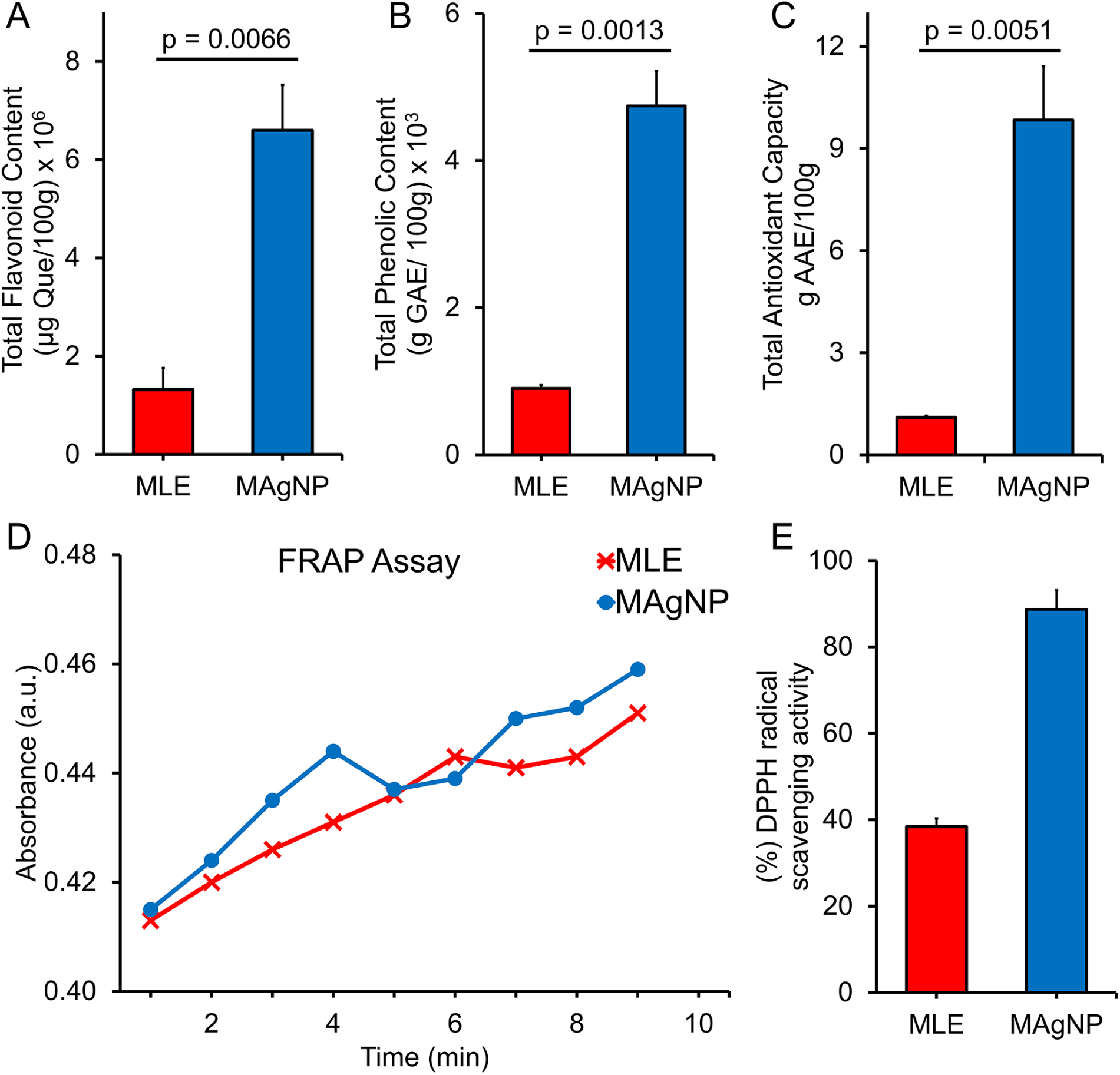
Antioxidant Assays with MAgNP. (A) Total Flavonoid Content of leaf extract and MAgNPs expressed as Quercetin equivalents (QE). MAgNPs showed much higher TFC values compared to the leaf extracts only. (B) Total Phenolic Content of leaf extract and MAgNPs expressed as Gallic acid equivalents (GAE). MAgNPs showed much higher TPC values compared to the leaf extracts. (C) Total Antioxidant Content of leaf extract and MAgNPs expressed as Ascorbic Acid Equivalents (AAE). MAgNPs showed much higher TAC values compared to the leaf extracts. (D) FRAP analysis of leaf extract and MAgNPs. The ferric ion reducing power was observed to be higher in MAgNPs compared to the leaf extracts. The results showed that MAgNPs exhibited greater free radical scavenging activity within 4 minutes and after 6 minutes, whereas the leaf extracts scavenging activity also increased but at a slower rate. (E) Percent DPPH values for leaf extracts and MAgNPs. The MAgNPs demonstrated superior free radical scavenging activity compared to the leaf extracts. The higher reducing power and electron-donating property of MAgNPs support the presence of a higher TFC demonstrated above. The error bars are the standard error of the mean (SE). P-values were obtained using a t-test to compare the means of the groups.

#### 3.8.2 Determination of Total Phenolic Content (TPC)

Total phenolic content was estimated as per the Folin-Ciocalteu (FC) colorimetric method described by Kumaresan *et al*. (Kumaresan et al., 2019). FC reagent is a mixture of tungstates and molybdates where its mechanism involves FC reagent being reduced in the presence of phenolic compounds in the plants, resulting in the formation of molybdenum–tungsten which resulted in blue-colored chromogen in an alkaline solution. Hence the technique is used to evaluate phenols with extreme absorption at 765 nm, the peak absorbance is proportional to the concentration of phenols (Blainski et al., 2013). MAgNPs showed higher TPC values compared to that of leaf aqueous extracts only (Figure 7 B), similar findings were found by Yarrappagaari, S. et al. also using green synthesized nanoparticles (Yarrappagaari et al., 2020).

#### 3.8.3 Determination of Total Antioxidant Content (TAC)

Antioxidant analysis was performed to quantify the ability of antioxidant compounds to quench free radicals. Most plant species possess phytochemicals such as phenols, flavonoids etc. that act as antioxidants, as reducing agents, or as single electron or proton donors for ROS and oxygen scavengers. Flavonoids are the most abundant class of polyphenolic compounds (Munteanu & Apetrei, 2021). Results showed that MAgNPs exhibit higher antioxidant values compared to that of leaf aqueous extract alone (Figure 7 C). Using green synthesized nanoparticles Patil and Raghavendra report similar results (S. Patil & Raghavendra, 2024).

Total Antioxidant Capacity was estimated as per the phosphomolybdenum assay (Perera & Kandiah, 2018). The principle implicates Mo (VI) is reduced to Mo (V) based on the electrons provided by the antioxidant in the sample for the formation of a green phosphate complex or Mo (V) (Saeed et al., 2012). Hence the technique is used to evaluate antioxidants with extreme absorption at 695 nm. Magnolia-synthesized silver nanoparticles showed higher TAC values compared to that of leaf extracts alone. Similar findings were found in Kandiah and Chandrasekaran (Kandiah & Chandrasekaran, 2021).

#### 3.8.4 Determination of Ferric Reducing Antioxidant Power (FRAP)

The FRAP assay evaluated the reducing power of leaf extract and MAgNPs. This involves the formation of a blue-colored compound by reduction of the Fe^3+^ tripyridyltriazine complex to Fe^2+^ tripyridyltriazine. The reaction is facilitated by electron donation from antioxidants at low pH (El Jemli et al., 2016). The results showed that MAgNPs exhibited a faster free radical scavenging activity, completing the process in just 4 minutes, whereas the leaf extract took longer (Figure 7 D). This indicates that MAgNPs possess a higher free radical scavenging activity than leaf extract alone. Ferric Reducing Antioxidant Power values were consistent with the Total Antioxidant (TAC) values reported elsewhere (Rajurkar & Hande, 2011).

#### 3.8.5 Determination of DPPH Activity

The DPPH assay was conducted to assess the free radical scavenging activity of MAgNPs compared to leaf extract alone. This method evaluates the electron-donating capacity of antioxidants, which neutralizes the DPPH radical by forming stable, diamagnetic molecules with an absorption maximum of 517 nm. The reaction involves a color change from purple/violet to colorless by the pairing of the odd electron on the nitrogen atom. The extent of decolorization indicates the magnitude of the reduction. The Beer-Lambert law was observed to hold within the range of absorptions (Noreen et al., 2017). Notably, MAgNPs demonstrated significantly higher free radical scavenging activity compared to the leaf extracts alone (Figure 7 E). These results are consistent with previous findings that MAgNPs have a higher TFC which generally correlates with free radical scavenging activity (Perera & Kandiah, 2018).

Results presented here suggest that MAgNPs have the potential to serve as an antioxidant therapeutic agent to address a variety of health conditions, such as cardiac arrest, Alzheimer’s disease, and diabetes, which are often linked to oxidative stress and inflammation. Future investigations should focus on elucidating the underlying mechanisms of MAgNPs’ antioxidant activity and exploring their potential applications in clinical settings.

### 3.9 Phytochemical Analysis

In the use of plant extracts in the synthesis of AgNPs, there are various bioactive compounds such as carbohydrates, proteins, saponins, steroids, tannins, and terpenoids which play a vital role in the synthesis, capping, bio-reduction, and stabilization of MAgNPs. These phytochemicals serve as reducing agents, donating electrons through functional groups like hydroxyl, carboxyl, and amine groups, which convert silver ions (Ag^+^) into elemental silver (Ag^0^). This process forms nanoparticles. Additionally, these compounds act as stabilizing agents, binding to the nanoparticle surface through hydrogen bonding or electrostatic attraction, preventing aggregation and controlling the size, shape, and stability of the nanoparticles. By capping the surface and preventing clumping, these phytochemicals ensure the formation of uniform and stable silver nanoparticles. Carbohydrate hydroxyl groups can reduce and stabilize nanoparticles by forming a surface coating. Protein amino acid residues can both reduce and stabilize nanoparticles through complex interactions. Saponins, with their amphiphilic nature, interact with both the aqueous phase and nanoparticle surface, providing stabilization. Steroids act as capping agents, creating a protective layer around nanoparticles. Tannins, polyphenolic compounds with multiple hydroxyl groups, readily reduce silver ions and stabilize nanoparticles. Terpenoids, with diverse structures and functional groups like hydroxyl and carbonyl, contribute to both the reduction and stabilization of silver nanoparticles (Jini et al., 2022).

### 3.10 Determination of Photocatalytic Activity

Photocatalytic activity refers to the ability of MAgNPs to facilitate a photoreaction under incident sunlight. When UV/visible light hits the surface of MAgNPs, it excites electrons from the conduction band (CB) to the valence band (VB), generating electron-hole pairs. The holes (H^+^) and electrons produced by VB and CB react with H_2_O and O_2_, respectively, forming hydroxyl radicals (OH•), superoxide ions (O_2_•-), and hydrogen peroxide radicals (HO_2_•). Together these radicals attack the azo bond in methyl orange, leading to dye degradation and the formation of intermediates which leads to the mineralization of the dye to a colorless final product (Figure 8 A) (Adam et al., 2018; Nagar & Devra, 2019). Green synthesized silver nanoparticles degrade methyl orange in the presence of sodium borohydride (NaBH_4_) by acting as a catalyst, facilitating the rapid electron transfer between the dye molecule and the reducing agent, effectively breaking down the dye structure and causing a significant decrease in its color intensity; this process is primarily driven by the large surface area and high catalytic activity of the silver nanoparticles, allowing for efficient degradation of the methyl orange dye molecule. This synergistic effect enables MAgNPs to efficiently degrade organic dyes within a short time. The absorption maximum of methyl orange at 460 nm decreased over time, indicating degradation (Figure 8 B). A shift of the methyl orange peak to 400 nm in the presence of MAgNPs was observed, suggesting a change in the molecular structure during degradation. Nagar and Devar proposed that the degradation pathway may involve the formation of intermediates through successive demethylation, leading to the substitution of the methyl group with hydrogen via homolytic cleavage of the nitrogen-carbon bond (Nagar & Devra, 2019). Similar findings were reported by Kandiah and Chandrasekaran (Kandiah & Chandrasekaran, 2021). This study demonstrates the potential of green synthesized nanoparticles to catalyze photoreactions for the degradation of non-biodegradable compounds that may contaminate natural water bodies, offering a promising approach to environmental remediation.

**Figure 8.**
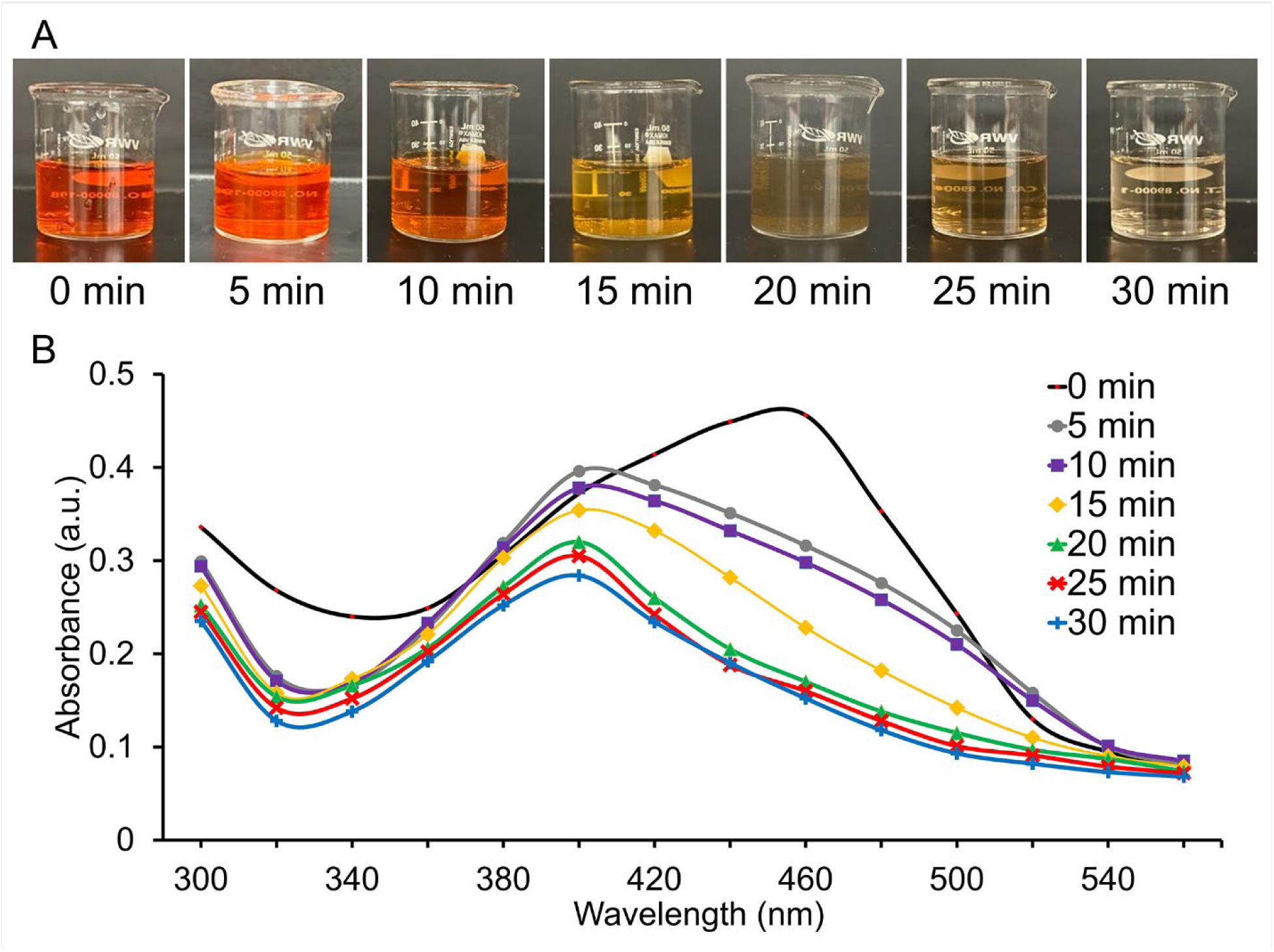
Photocatalytic activity of MAgNP on methyl orange. Radicals generated by exposure to sunlight attack the azo bond in methyl orange leading to the degradation of the dye and formation of intermediates, ultimately resulting in the mineralization of the dye to a colorless final product in 30 minutes.

## 4. Conclusion

In conclusion, green synthesis of MAgNPs is rapid, cost-effective, and reproducible. Silver nanoparticles, particularly when synthesized from natural sources like Magnolia, offer broad applications in medicine, antioxidant therapies and pollution control. Their bioactive properties and environmental compatibility suggest great potential in the quest for safer, more effective solutions across various fields. Magnolia-synthesized silver nanoparticles have clearly been shown to have both antibacterial and antifungal properties against known bacterial and fungal species of medical interest thus rivaling costlier chemical antimicrobials currently in use. The ability to inhibit bacteriophage activity can serve as a model for potential applications against human viruses, positioning MAgNPs as promising candidates for antiviral drug development.

The dual anticancer and antimicrobial properties of MAgNPs suggest potential applications in cancer therapy where secondary infections are a concern. The ability to simultaneously target cancer cells and prevent bacterial colonization could be particularly valuable in immunocompromised cancer patients. Magnolia silver nanoparticles have the potential to serve as an antioxidant therapeutic agent in the treatment of a variety of free radical-induced disorders. This study also demonstrates the potential of MAgNPs to catalyze photoreactions for the degradation of non-biodegradable compounds, offering a promising approach to environmental remediation. We are continuing to explore the green synthesis of nanoparticles using other metals, other plants, and plant parts and their potential uses in targeted nanomedicine.

## Abbreviations

AgNPs: silver nanoparticles
MLE: Magnolia Leaf Extract
MAgNPs: Magnolia silver nanoparticles
ROS: Reactive oxygen species
TFC: Total Flavonoid Content
TPC: Total Phenolic Content
TAC: Total Antioxidant Capacity
DPPH: 2, 2-Diphenyl-1-picrylhydrazyl
FRAP: Ferric Reducing Antioxidant Power
SEM: Scanning Electron Microscope
FC: Folin-Ciocalteu
MRSA: Methicillin Resistant *Staphylococcus aureus,*
MSSA: Methicillin Sensitive *Staphylococcus aureus*
QE: Quercetin equivalents
GAE: Gallic acid equivalents
AAE: Ascorbic acid equivalents
ZOI: Zone of Inhibition
RT: Room temperature

## CRediT authorship contribution statement

Shaveen De Mel: Conceptualization and Designing the Research Project, Writing, Performing Experiments, Visualization, Validation, Methodology, Investigation, and Formal Analysis. Juliana Gruenler, Logan Khoury, Ashton Heynes: Graphical Abstract and Laboratory Bench Work. Julianne Frazekas, Kendra Damaske: Laboratory Bench Work. Thushara Galbadage: Writing Methodology Results and Discussion, Visualization, Conceptualization, Editing, and Software. Richard S. Gunasekera: Investigation, Methodology, Conceptualization, Funding Acquisition, Writing and Reviewing, Visualization, Supervision. Ross S. Anderson: Supervision, Investigation, Methodology, Conceptualization, Writing, Reviewing, and Editing.

## Acknowledgments

The authors express gratitude to The Master’s University (TMU) & Biola University (BU) for granting resources and financially supporting this research. We thank Dr. Joe Francis, Professor Michael Kornoff, Thai Perez, David Fernando, Shaina Job, and Natalia Soto from TMU & Randil Jayasekara from BU for their contributions in the preliminary stage of the research. We wish to convey our gratitude to Shavindri De Mel for data analysis and graphics. We also wish to thank Drs. Sarah Maithel and Kevin Nick, Department of Earth and Biological Sciences, Loma Linda University for use of the SEM.

